# Comparison of different head and neck positions and behaviour in ridden elite dressage horses between warm-up and competition

**DOI:** 10.1101/2021.12.17.473217

**Authors:** Kathrin Kienapfel, Lara Piccolo, Annik Gmel, Dominik Rueß, Iris Bachmann

**Author notes:** Corresponding author: Kathrin Kienapfel.

## Abstract

Head and neck position (HNP) has been identified in literature as important influence on wellbeing. It was investigated in ridden elite dressage horses whether there is a relation between the HNP, ethological indicators and the grading in the warm-up area and in the test. 49 starters (83%) of a Grand-Prix Special (CDIO5 *) as part of the CHIO in Aachen 2018 and 2019 were examined. For each horse-rider pair, HNP (angle at vertical (AT), poll angle (PA), neck angel (NA)) used were analysed as well as conflict behavior for 3 minutes each in warm-up area and test. 6571 individual frames were used. The noseline was carried significantly less behind the vertical in test vs. warm-up (5.43 ° ± 4.19 vs. 11.01 ° ± 4.54 behind the vertical; T = 34.0; p < 0.05). The horses showed significantly less conflict behavior in the test vs. warm-up (123 ± 54 vs. 160 ± 75) (T = 76.00; p < 0.01). In the latter, a smaller PA and more defensive behaviour of the horses was observed compared to the test. A correlation between the grading of test and HNP was found (R = 0.38; p < 0.05). The further the noseline was behind the vertical, the higher was the chance of a good rating. The higher riders were ranked in the “FEI world ranking”, the higher were their marks in the competition (2018: r = -0.69, p < 0.05; 2019: r = -0.76, p < 0.05). Horses of riders higher in world ranking tended to show more unusual oral behaviour (r = -0.30, p < 0.05), and a noseline stronger behind the vertical (r = - 0.37, p < 0.05) resulting in a smaller NA (r = 0.43, p < 0.05). This are from the point of view of animal welfare problematic results.

## Introduction

In recent years, horse riding at an elite level has come under increased scrutiny in relation to animal welfare. In the Olympic discipline of dressage, whether riding in certain HNPs (head-and-neck positions) may compromise the welfare of horses has been heatedly discussed. Official federations regularly state the importance of a sound and happy athlete (Federation Equestre Internationale, 2019), while in practice, equine welfare is often overlooked when the horse is performing well. In numerous studies, the influence of HNPs on different parameters of health and welfare of horses was evaluated ([1-9]. In nearly all of these studies, the hyperflexed position stood out as being extreme. The consequences of this HNP were distress and compromised welfare evidenced by physiological constraints seen in increased heart rate (HR), cortisol levels and conflict behaviours or decreased heart-rate variability (HRV) [5, 10-14]. In addition, some studies detected impaired breathing [15-17] and potentially pathological occurences in the neck [4, 18-22], while being ridden with the noseline behind the vertical. In 2015 a metaanalysis quantified and analysed the assembled scientific results and found a significantly negative influence of HNPs with the noseline behind the vertical on horse welfare [23].

In elite dressage competitions, the directive is clearly defined. According to the FEI dressage rules the noseline has to be at or slightly in front of the vertical at all times [24]. Hyperflexion as a form of aggressive riding is forbidden, while not being defined in more than the most excessive form where the chin of the horse is almost touching its chest [25] (https://www.youtube.com/watch?v=2tpVNSTR8Nw). If riders start in competitions they should self-evidently follow these rules to receive high grades and maintain the welfare of the horse. If the ideal HNP is not achieved constantly at all times (eg, while warming up) it should of course be the final goal. Therefore, we chose world class riders to assure the best possible research results of skilled and experienced horse-rider-pairs. We expected to find the whole range of HNPs, knowing that from discussions of the last 10 years the ideal is not always maintained. However, following the FEI statement “the welfare of the horse is paramount”, the occurrence of Hyperflexion is expected as seldom event, with desirable riding to be present most of the times in elite sports [26].

In the present study the recent situation in dressage riding of elite horses was evaluated in two selected typical competitions. The world leading elite was chosen to provide the best possible horse-rider relationship and skill of the rider. Additionally, at this level, equine health and management are at very high standards. In the context of elite competitions, the horses are checked by veterinarians each time as “fit to compete” so this study only included sound horses. The aim of the study was the evaluation of the relationship of HNP, behavioural parameters and the marks given by the judges for the competition run. Furthermore, the relationship between the HNP used in the warm-up area and the competition was studied.

## Methods

For the study, the used head-neck positions and behaviours indicative of conflict of 49 starters of a Grand Prix special (CDIO5*) of the CHIO in Aachen were investigated. 42% (n=22) of the starters were in the “Top 20” of the world during the respective year of the data collection, and 86% in the “Top 100”, resulting in a sample of the best riders in the world. In 2018, 73% and in 2019 93% of the starters were evaluated in both situations: warm-up and competition. In total, 90% of all starters of the Grand Prix special were included in the study. For each horse-rider pair, the head-neck positions as well as the conflict behaviors (Table 1) were analysed for three minutes each in the warm-up area and during the competition. Ear position and movement were analysed in a separate study (in prep). Filming was done with either a Sony FDR-AX 53 or a Sony HDR-CX625 in HD-quality. The researcher team recorded the videos of the riders in the warm-up area whereas the videos of the competition were provided by an Internet video platform. Each video was cut in sections of continuous riding of three consecutive minutes if available after the beginning of the working phase (defined as starting of sitting trot). Phases of walk, standing or other horses crossing in front of the observed pair were cut out. No other selection was done. Every single frame of the videos with visible horses’ profile view was analysed by a newly developed annotation tool (Group ‘Computer Vision’, Institute of Informatics, Humboldt University Berlin). Four anatomical markers (mouth, neck, shoulder and withers) were set by hand in the annotation tool. With these anatomical markers, three angles were determined: The angle of the nose line in relation to the vertical (α), the poll angle (β) and the angle between the shoulder and the withers (γ) (fig 1). An angle in front of the vertical from the horses viewpoint was defined as all angles α<0, behind the vertical as α>0, meaning the angles behind the vertical were given as positive values.

**Table 1:**

| Behaviour | Explanation |
| --- | --- |
| Rearing | The horse’s forebody and forelimbs rise, such that the weight is borne by the hind limbs while standing. Every leaving of the ground was counted as one event. |
| Bucking | The horse lowers the head and neck and raises the hindlimbs off the ground while standing. Every act is counted as one event. While moving, see “errors in rhythm”. |
| Unusual oral behaviour (all deviations from a still and closed mouth or chewing with closed lips) | The horse opens the mouth so a gap between the upper and lower jaw is visible, showing the teeth or tongue for more than 1 s, chewing movements with visible separation of upper and lower jaw or moving the lower jaw opposite to upper jaw while chewing. The total time of these behaviours was taken and counted as one event per second. |
| Tail swishing | The horse moves the tail in a fast motion in a vertical, horizontal or combined direction. Every act was counted as one event independent of the vigor of the movement. |
| Headshaking | The horse moves the head quickly up and downwards and/or from side-to-side. Every act was counted as one event. |
| Crabbing | Hind limbs of the horse do not follow the track of the front limbs. The total time of this behaviour was taken and counted as one event per second. |
| Errors in rhythm | Insertion of an additional step resulting in loss of the rhythm. Bucking (throwing the hindquarters up simultaneously), kicking with one leg and every other hopping movement outside the specific gait while |
|  | moving was included here. Every deviation from the gait specific rhythm was counted as one event. |
| Nose tilting | The horse tilts its nose to one side. The duration of this action was taken and counted as one event per second |
| Going-against-reins | The horse pulls against the reins and breaks the line between elbow of the rider and rings of the bit. Every act was counted as one event. |
| Ear movement | analysed in other study |
| Ear position | analysed in other study |

**Fig. 1:**
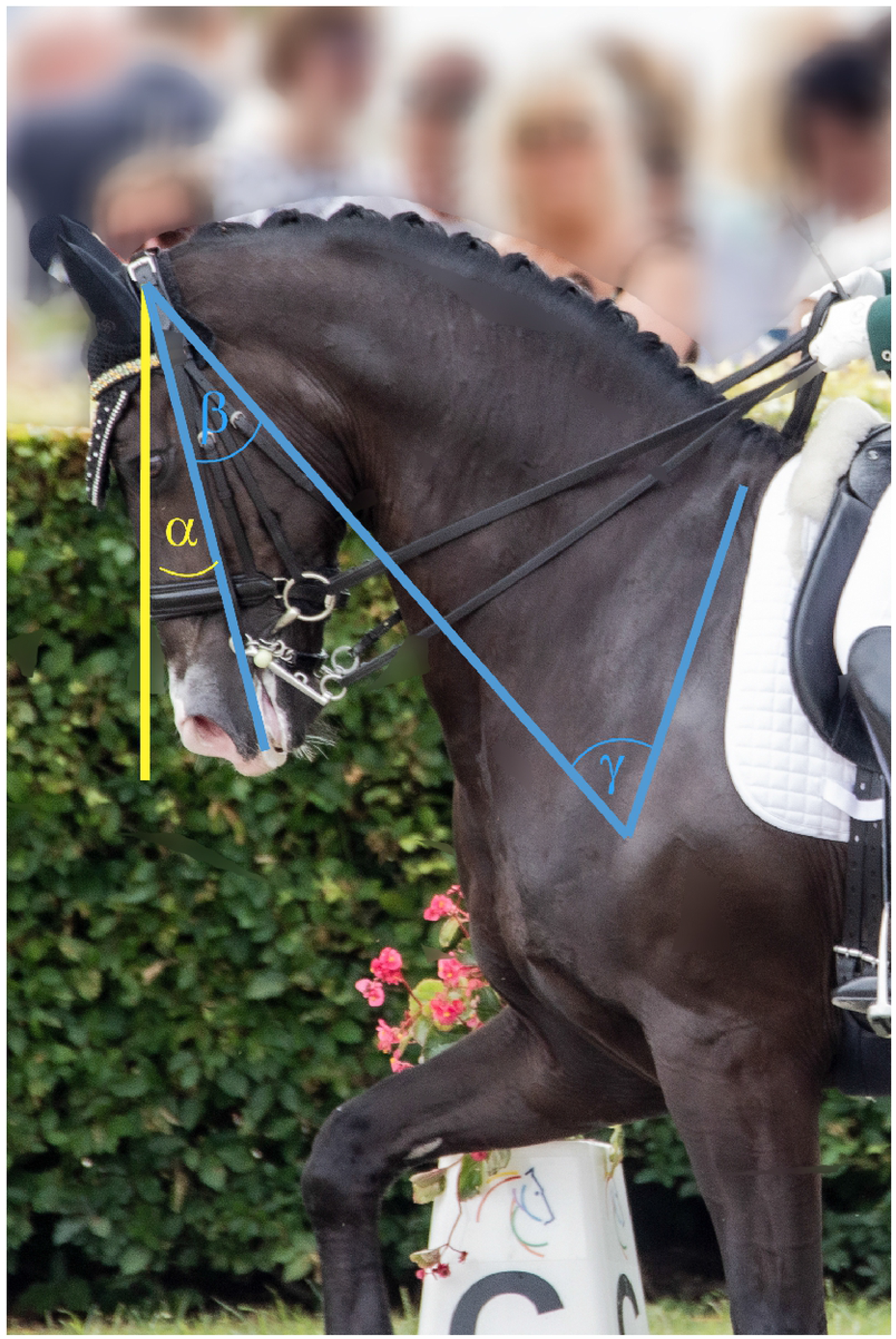
Studied angles: α= angle at the vertical, β= poll angle, γ= shoulder angle (© Kienapfel)

For each horse, 33 ± 12 single frames in the warm-up area and 103 ± 24 single frames in the competition were analysed (total: 6,571 individual frames, Table 2). All behaviours indicative of conflict were analysed in the Observer XT (Noldus) using the focus animal method. Every occuring behaviour was noted following the ethogram (Table 1). The data was not normally distributed (Shapiro Wilk test, p > 0.05). General linear models were created to determine the different influences on the parameters. In addition, Wilcoxon tests were performed. Furthermore, the parameters “age”, “gender” and “breed” of the horses, as well as the “Competition Ranking” and “World Ranking” were tested for correlations using Spearman Rank Correlation Test.

**Tab. 2:** Number of analysed single frames

| *Year* | | Absolute numbers of frames | Frames per horse (± standard deviation) |
| --- | --- | --- | --- |
| *2018* | Warm-up | 736 | 37±15 |
| | Competition | 2007 | 96±24 |
| *2019* | Warm-up | 750 | 28 (± 10) |
| | Competition | 3078 | 109 (±24) |
| ***Total*** |  | **6571** | **135 (±18)** |

The distribution of the final scores from the five judges were visualised in violin plots. The interrater reliability between the scores was estimated using intra-class correlation coefficients with the r package irr (Gamer et al. 2021).

## Results

During warm-up and competition, the horses showed different amounts of several conflict behaviours (Table 3).

**Table 3:**
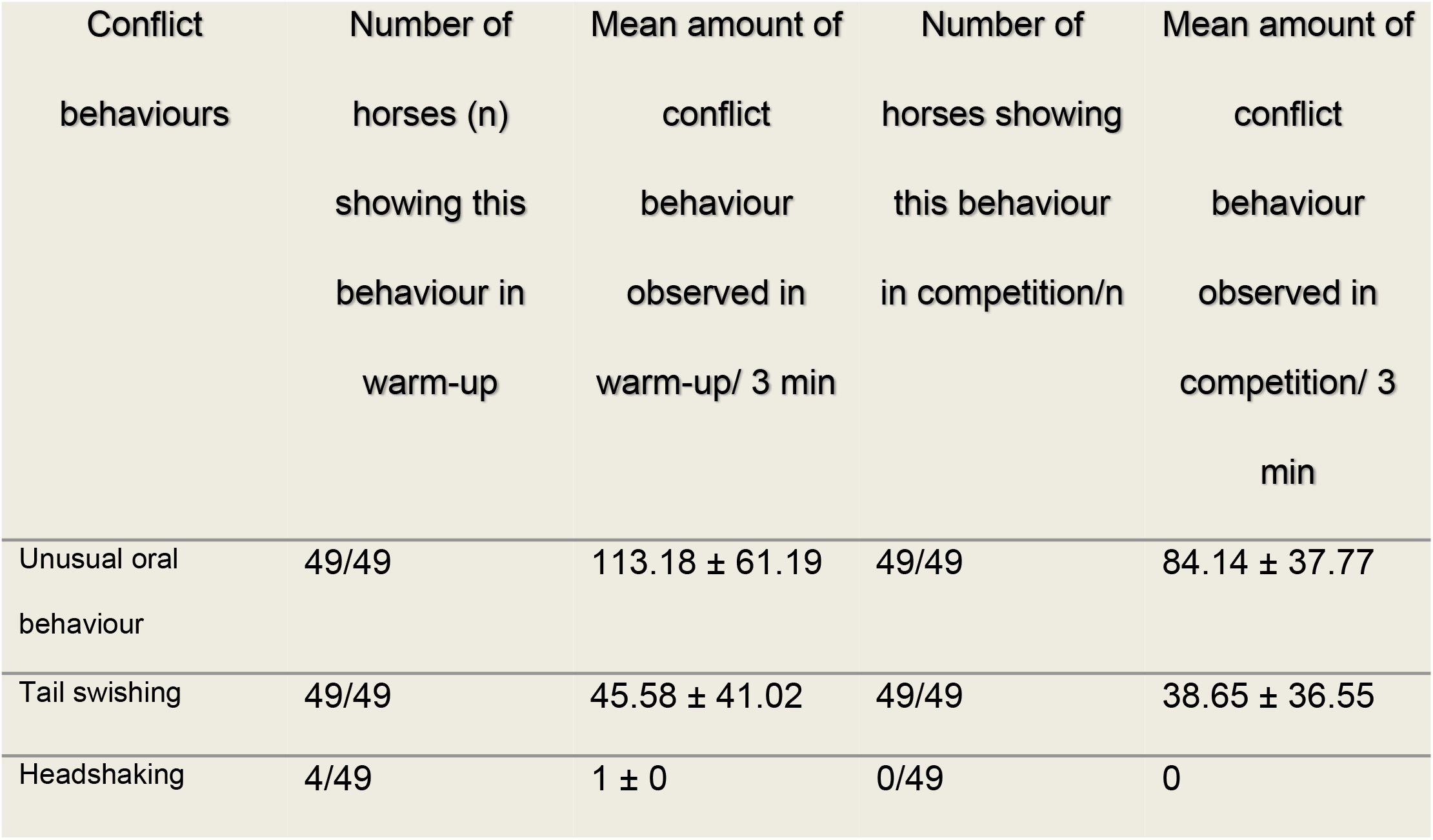
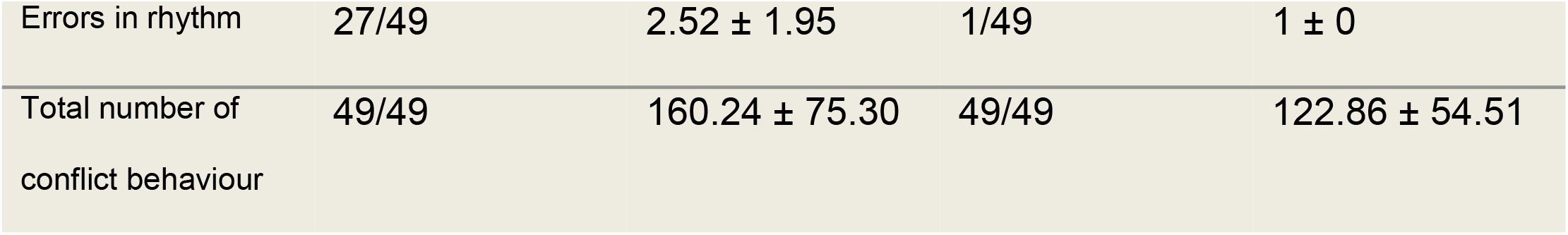
Mean amounts of conflict behaviours observed during warm-up and competition in three minutes (only observed behaviour categories are listed)

| Conflict behaviours | Number of horses (n) showing this behaviour in warm-up | Mean amount of conflict behaviour observed in warm-up/ 3 min | Number of horses showing this behaviour in competition/n | Mean amount of conflict behaviour observed in competition/ 3 min |
| --- | --- | --- | --- | --- |
| Unusual oral behaviour | 49/49 | 113.18 ± 61.19 | 49/49 | 84.14 ± 37.77 |
| Tail swishing | 49/49 | 45.58 ± 41.02 | 49/49 | 38.65 ± 36.55 |
| Headshaking | 4/49 | 1 ± 0 | 0/49 | 0 |
| Errors in rhythm | 27/49 | 2.52 ± 1.95 | 1/49 | 1 ± 0 |
| Total number of conflict behaviour | 49/49 | 160.24 ± 75.30 | 49/49 | 122.86 ± 54.51 |

Significant differences of the head-neck position and behavioural parameters were found between the warm-up and the competition situation. The horses’ nasal plane was more behind the vertical during the warm-up than during the competition (angle α: 11.01° ± 4.54° vs. 5.42° ± 4.19°; T = 34.0; p < 0.05, Figure 2). The poll angle (β) was significantly higher during the competition than during the warm-up (27.8° ± 3.56° vs. 23.42° ± 3.07°, T = 63.00; p < 0.05). There was no significant difference of the angle γ between these two situations (p > 0.05). Furthermore, not only the head-neck position, but also the amount of conflict behaviour differed during the warm-up and the competition. The horses showed significantly more conflict behaviour during the warm-up than during the competition (160.24 ± 75.30 vs. 122.86 ± 54.51; T = 261.00; p < 0.05, figure 3). The amount of tail swishes was not significantly different during the warm-up and the competition (45.58 ± 41.02 vs. 38.65 ± 36.55). A significant difference of the amount of unusual oral behaviour between the warm-up and the competition was ascertained (113.18 ± 61.19 vs. 84.14 ± 37.77; T = 327.00; p < 0.05). Errors in rhythm (2.52 ± 1.95 vs. 1 ± 0) and headshaking (1 ± 0 vs. 0) were rarely observed, especially during competition in only a few horses.

**Fig. 2:**
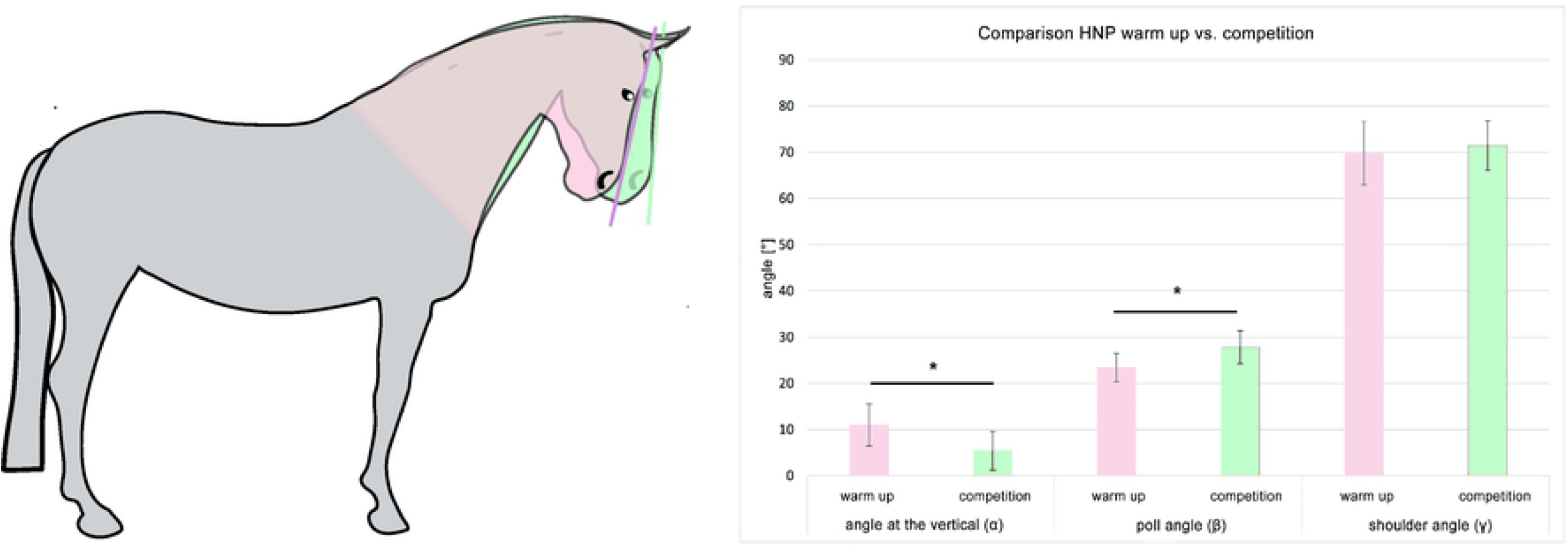
Angles of the head and poll in warm-up area and competition © Kienapfel

**Fig. 3:**
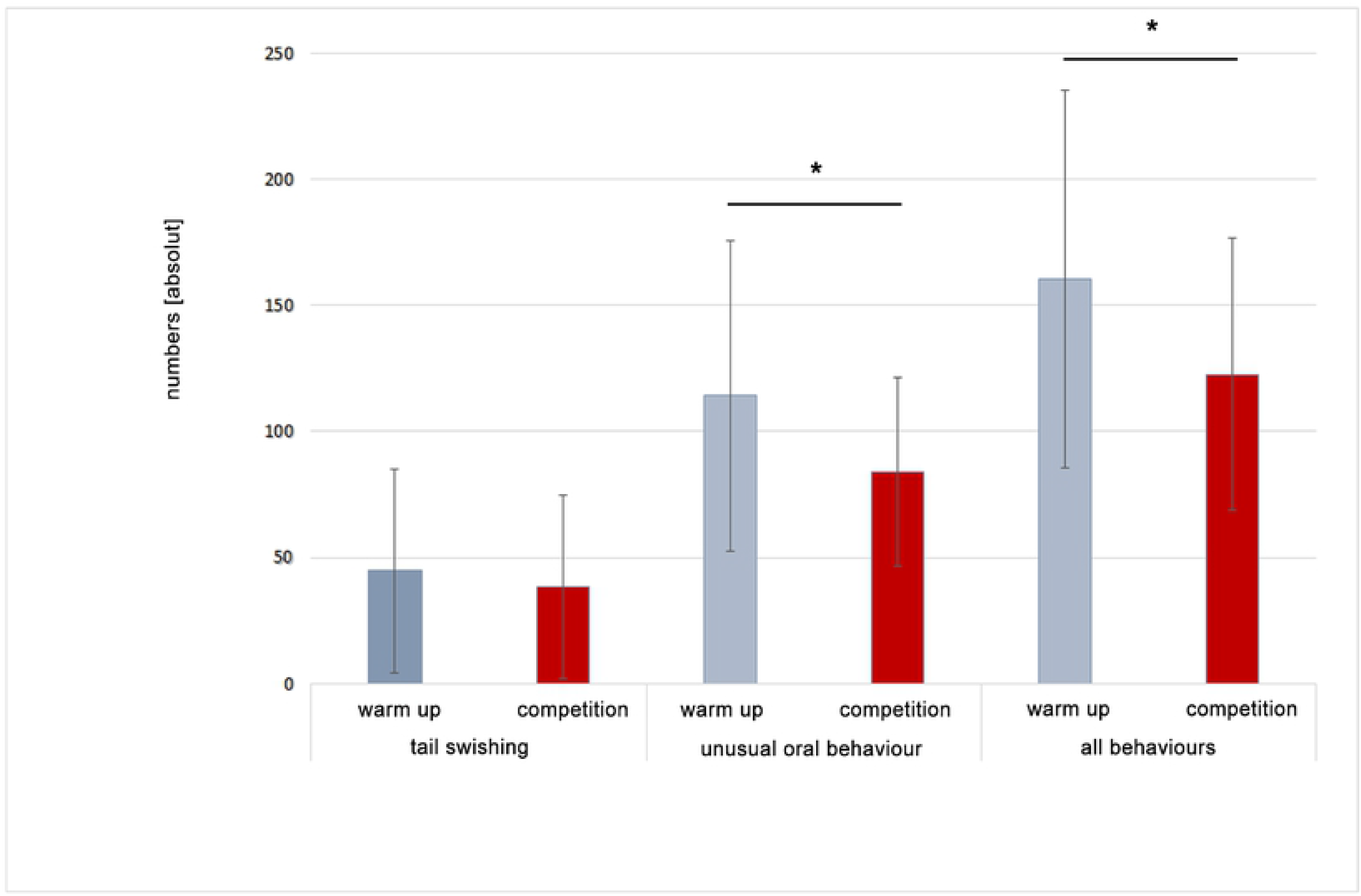
Ethological parameters warm-up vs. competition © Kienapfel

Additionally, the concordance of the five judges’ scores during the competitions was investigated. The judges’ scores correlated strongly among each other (R > 0.96, p < 0.001, figure 4). The ICC was excellent (ICC=0.99, CI=0.98-0.99). Furthermore, the scores were positively correlated with the angle α (R = 0.38, p < 0.05). The more the horses’ nasal plane was behind the vertical, the higher was the chance for a higher score. Age, breed or gender had no influence on the analysed parameters.

**Figure 4:**
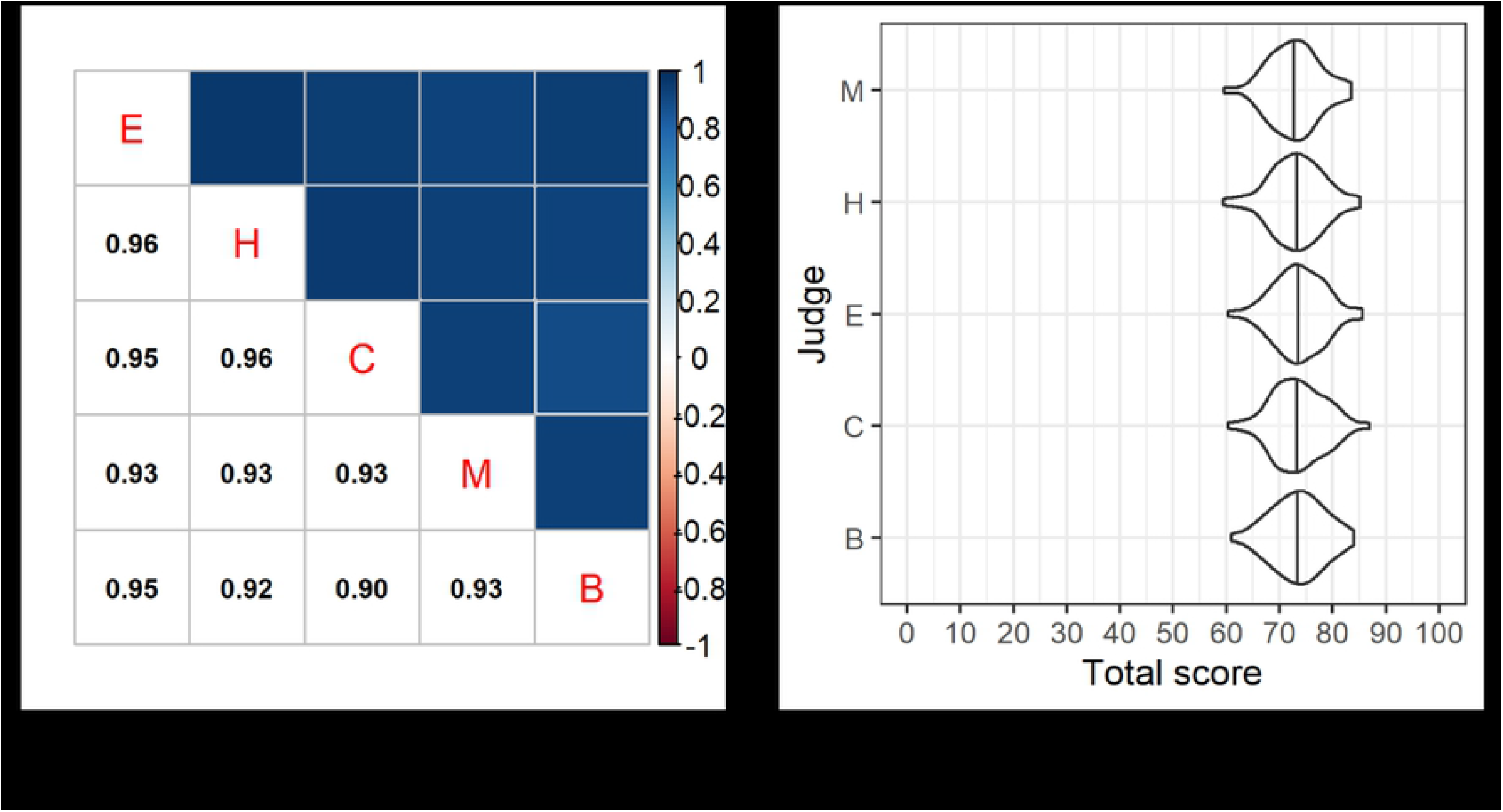
(A) Cross-correlation matrix of the judges for the total score, pooled for both years by position (E, H, C, M and B). (B) Violin plots of the scores given by each judge, pooled by position (E, H, C, M and B)

The parameter “World Ranking” was strongly correlated with the “Competition Ranking” in both years (2018: r = -0.69, p < 0.05; 2019: r = -0.76, p < 0.05), meaning the higher the riders were ranked in the world ranking, the higher were their marks in the competition.

Furthermore, the “World Ranking’’ parameter was also correlated with the amount of unusual behaviour (r = -0.30, p < 0.05), with the angle of the noseline behind the vertical (r = -0.37, p < 0.05) and the poll angle (r = 0.43, p < 0.05). So the horses of riders higher in the world ranking, tended to show more unusual oral behaviour and a noseline more behind the vertical resulting in a smaller poll angle.

### Discussion

A majority of the starters (83%) of one of the most important international competitions were examined in this study for 2 years in a row, providing an insight into the current international dressage sport. The HNP was analysed by an annotation tool custom made for this project. This is a very precise method of analysing biomechanical parameters without attaching markers on the animals directly, yet allowing the analysis of a large number of single frames. In total, the three angles were determined for the specification of the HNP used in each case for 6,571 individual frames, evenly distributed over the respective sequences, maximising an independence of gait-related variations. Even so, it would be better to have continuous angle detection to assure full independence of gait and perspective, but that is not possible with current technical instruments in a field situation and additionally here by looking at high level competitions we could not attach any sensors to the horses. Further technical improvements would be very welcome.

Significant differences were found between riding in the warm-up area and in the competition. During the competition, horses showed significantly less conflict behaviour than in the warmup area. [5] already found similar results by comparing these two situations in horses that started at a significantly lower level in small national competitions in Germany.

A vast majority of conflict behaviour was tail swishing and unusual oral behaviour. No difference in tail swishing could be found in the two studied conditions (45.58 ± 41.02 vs. 38.65 ± 36.55). Although the HNP was changed in the test, the level of tension in horse and rider might increase in the competition situation, where the performance should be the best possible as the final performance is being judged. This points to a possible correlation of other factors causally influencing tail swishing more than HNP alone. In addition, sequences of tasks like piaffe, passage and pirouettes are demonstrated in the test situation more often than in the warm-up, possibly resulting in greater physical stress. However, frequent tail swishing is generally seen as an alarm signal while riding [5, 10, 18, 28]. Further investigations are therefore required to determine the influence of specific ridden tasks on the amount of tail swishing in elite dressage horses.

The amount of unusual oral behaviour differed significantly between both analysed situations and the same was true for the used HNPs. In the competition, horses were presented with larger neck angles and therefore a noseline less strongly behind the vertical. The unusual oral behavior seems to be closely related to the head and neck position and could be a useful indication for the rider to acknowledge a necessary change in the selected HNP. It should be emphasised that only one horse in the sample was ridden with the noseline in front of the vertical - therefore, for the sake of clarity, the angles behind the vertical were given as positive values.

Another significant connection in this study is the relationship between the judges’ rating and the HNP. None of the other examined parameters correlated with the grade (age, race and behavior). The stronger the horses with the noseline behind the vertical in the test, the higher the probability of a good placement (R = 0.38, p < 0.05). This is surprising insofar that the national and international regulations require an HNP with the noseline at or slightly in front of the vertical [24]. Even more so, as, according to a study, the judges tend to focus on the forehand (head, neck and shoulder area) of the horse [29], so this relatively easy visible indicator should be taken into account. The extremely high ICC (0.99) in this study was consistent with findings that the agreement between judges scoring elite competitions, with better-known riders and horses, was higher than when scoring novice [30]. This may be due to the judges having higher qualifications at the elite level, an implicit bias for specific, easily recognizable rider-horse pairs, or a combination of both. The high scores up to 87% were also reflective of the word ranking level of the competitions considered in this study, as the highest scores are awarded to the best rider-horse pairs in dressage. The scores were given with a low variance of scores between 59 and 87% with possible scoring of 0-100%. This relatively narrow range of the regular score may be a reason for the high ICC. To gain a more complex image of the performance of each-horse-rider pair it could be beneficial to take advantage of the full scoring range following the definitions in the rulebooks.

However, high agreement does not necessarily indicate validity, as the higher scores were positively correlated to HNP behind the vertical, which should in theory be penalised. Apparently the reason for this undesirable correlation indicates the failing of following own intern rules, which has to be based on other factors than agreement. This study revealed a discrepancy between FEI rules and scores in relation to objectively measured kinematic and behavioural parameters, which needs to be addressed in order to improve equine welfare at the highest levels of the sport.

